# Quantifying synergy in the bioassay-guided fractionation of natural product extracts

**DOI:** 10.1101/2020.06.23.166686

**Authors:** Micah Dettweiler, Lewis Marquez, Max Bao, Cassandra L. Quave

## Abstract

Mixtures of drugs often have greater therapeutic value than any of their constituent drugs alone, and such combination therapies are widely used to treat diseases such as cancer, malaria, and viral infections. However, developing useful drug mixtures is challenging due to complex interactions between drugs. Natural substances can be fruitful sources of useful drug mixtures because secondary metabolites produced by living organisms do not often act in isolation *in vivo*. In order to facilitate the study of interactions within natural substances, a new analytical method to quantify interactions using data generated in the process of bioassay-guided fractionation is presented here: the extract fractional inhibitory concentration index (EFICI). The EFICI method uses the framework of Loewe additivity to calculate fractional inhibitory concentration values by which interactions can be determined for any combination of fractions that make up a parent extract. The EFICI method was applied to data on the bioassay-guided fractionation of *Lechea mucronata* and *Schinus terebinthifolia* for growth inhibition of the pathogenic bacterium *Acinetobacter baumannii*. The *L. mucronata* extract contained synergistic interactions (EFICI = 0.4181) and the *S. terebinthifolia* extract was non-interactive overall (EFICI = 0.9129). Quantifying interactions in the bioassay-guided fractionation of natural substances does not require additional experiments and can be useful to guide the experimental process and to support the development of standardized extracts as botanical drugs.

## Introduction

The study of combination therapy—the treatment of disease with mixtures of drugs—is a growing field in modern pharmacology, but an old science as far as the actual application of medicine is concerned (1). While single-compound drugs have revolutionized the treatment of many conditions, the development of resistance, among other factors, has prompted a return to combination therapies for several diseases, including cancer (2), malaria (3), HIV (4), and antibiotic-resistant infections (5). Combination therapy has a variety of potential benefits, including synergy: greater potency of a drug mixture than would be predicted from the activity of each constituent drug. Drugs that interact synergistically in a mixture have the advantage of requiring lower doses for efficacy compared to their isolated use (6), but even in the absence of synergy, drug mixtures can exhibit lower toxicity and slow the development of resistance (7).

A major obstacle to the development of combination therapies is the complexity that exists by definition in mixtures (8). Determining efficacy, toxicity, and pharmacokinetics is difficult enough with a single compound, and each additional factor multiplies the study needed to vet a medicine properly. Additionally, identifying which drugs to combine and in what ratio can be arbitrary and very time-consuming, particularly when investigating mixtures of more than two drugs. Finally, there has been a struggle to develop and agree upon valid models, assays, and analysis of the interactions present in drug mixtures (9, 10).

One potential pathway to identifying useful combination therapies for human health is investigating natural substances. This heuristic may be productive because the mixtures of compounds produced by living organisms are not arbitrary; natural selection may favor organisms that produce a secondary metabolite profile that contains profitable interactions between compounds as well as compounds that are active by themselves (8, 11). The investigation of natural substances for combination therapies seeks to draw on this elaborate chemistry to identify pre-existing drug mixtures with useful bioactivity.

The advantages of complex natural substances in medicine are not simply theoretical. Many modern drugs have been derived from or inspired by natural substances used by humans for medicine, such as digoxin from *Digitalis* spp. and morphine from *Papaver somniferum* (1). Moreover, there are many examples of crude natural substance extracts exhibiting greater activity than would be predicted from their most active component, as in the case of *Artemisia annua* and artemisinin (12). It is worthwhile, then, in the search for new combination therapies, to consider natural substances.

Previous investigations of synergistic activity in natural substances have largely focused on pairwise combinations of natural compounds with existing medications or two natural compounds with each other (13), and while these experiments provide useful information, it is impossible to optimally use natural substances as medicines without understanding intra-extract interactions on a broader scale. It is acknowledged in the literature that synergy often exists in extracts (8), but only rudimentary steps have been taken to describe it when it occurs (14). Using the basic equations of Loewe additivity (15), however, it is possible to both identify and quantify interactions in combinations of any number of drugs (16).

In the process of bioassay-guided fractionation, an extract is tested for activity, then chemically separated, then the resulting fractions are tested for activity (14). This is an iterative process; the most active fraction can itself be separated and have its fractions tested, repeating until a single or multiple active compounds are isolated. Bioassay-guided fractionation creates a reversed approach to drug interaction analysis; instead of creating and testing a mixture, a naturally existing mixture is separated and tested such that interactions present in the mixture are elucidated. By treating natural substance extracts as combinations of their fractions, data collected from bioassay-guided fractionation can be used to quantify synergy in bioactive extracts without additional experiments and inform go/no go decisions. When activity is lost in the process of bioassay-guided fractionation, quantified synergy can support the pursuit of alternate drug development directions such as the FDA Botanical Drug Pathway (1).

The objective of this study is to present the extract fractional inhibitory concentration index (EFICI), a new application of existing analytical methods, for use in the quantification of synergy and other interactions in natural substances. To demonstrate this method, we selected two plant species used as anti-infectives in traditional medicine, *Lechea mucronata* Raf. (Cistaceae) (hairy pinweed) and *Schinus terebinthifolia* Raddi (Anacardiaceae) (Brazilian peppertree) (17), and subjected their extracts to bioassay-guided fractionation for growth inhibition of *Acinetobacter baumannii*, a clinically relevant pathogen (18). EFICI analysis was performed for both extracts.

## Materials and Methods

### Bioassay-guided fractionation of *L. mucronata* and *S. terebinthifolia*

*Lechea mucronata* roots were collected with permission at the Jones Center at Ichauway ecological field station in June 2018 in Baker County, Georgia, USA. *Schinus terebinthifolia* leaves were collected in November 2017 in DeSoto County, Florida, USA on private lands with permission from the landowners. Plant identity was confirmed by Dr. Cassandra Quave, and voucher specimens (CQ-793 and CQ-651) were deposited in the Emory University Herbarium (GEO), available for viewing online via the SERNEC portal (19). The roots of *L. mucronata* (CQ-793) and the leaves of *S. terebinthifolia* (CQ-651) were dried in a dehumidified cabinet, ground to a fine powder using a Wiley Mill Plant Grinder (0.5 mm mesh) and extracted via two 80% ethanol (v/v) macerations (1:10 w/v), each for 72 hours at room temperature. The extracts, named 1835 (*L. mucronata* roots) and 429 (*S. terebinthifolia* leaves), were vacuum filtered, concentrated *in vacuo*, and stored at −20° C until further use.

Extract 1835 (*L. mucronata*) was partitioned with a modified Kupchan liquid-liquid partitioning scheme, yielding hexane, ethyl acetate, and water partitions named B, C, and F, respectively. The water partition (1835F) exhibited the greatest growth inhibition against *A. baumannii* and underwent further fractionation through reversed-phase high-performance liquid chromatography (HPLC). The HPLC method development was performed using an Agilent 1260 Infinity analytical HPLC system with an XDB-C18 4.6×250 mm, 5 μm column. A two-solvent gradient system starting from a 98:2 mixture of 0.1% formic acid in water and 0.1% formic acid in methanol exhibited the best separation when monitored at 254 nm over 100 minutes. The analytical method was subsequently adapted for preparative-HPLC by taking into account the differences in column size and flow rate. Preparative HPLC was performed using an Agilent 1260 Infinity II system with an XDB-C18 30×250mm, 5μm column. Two sequential runs were performed, each with a 5.0 mL sample injection (23 mg/mL in 80:20 H_2_O:MeOH), and a total of 22 fractions were collected (Fig 1).

**Figure 1.**
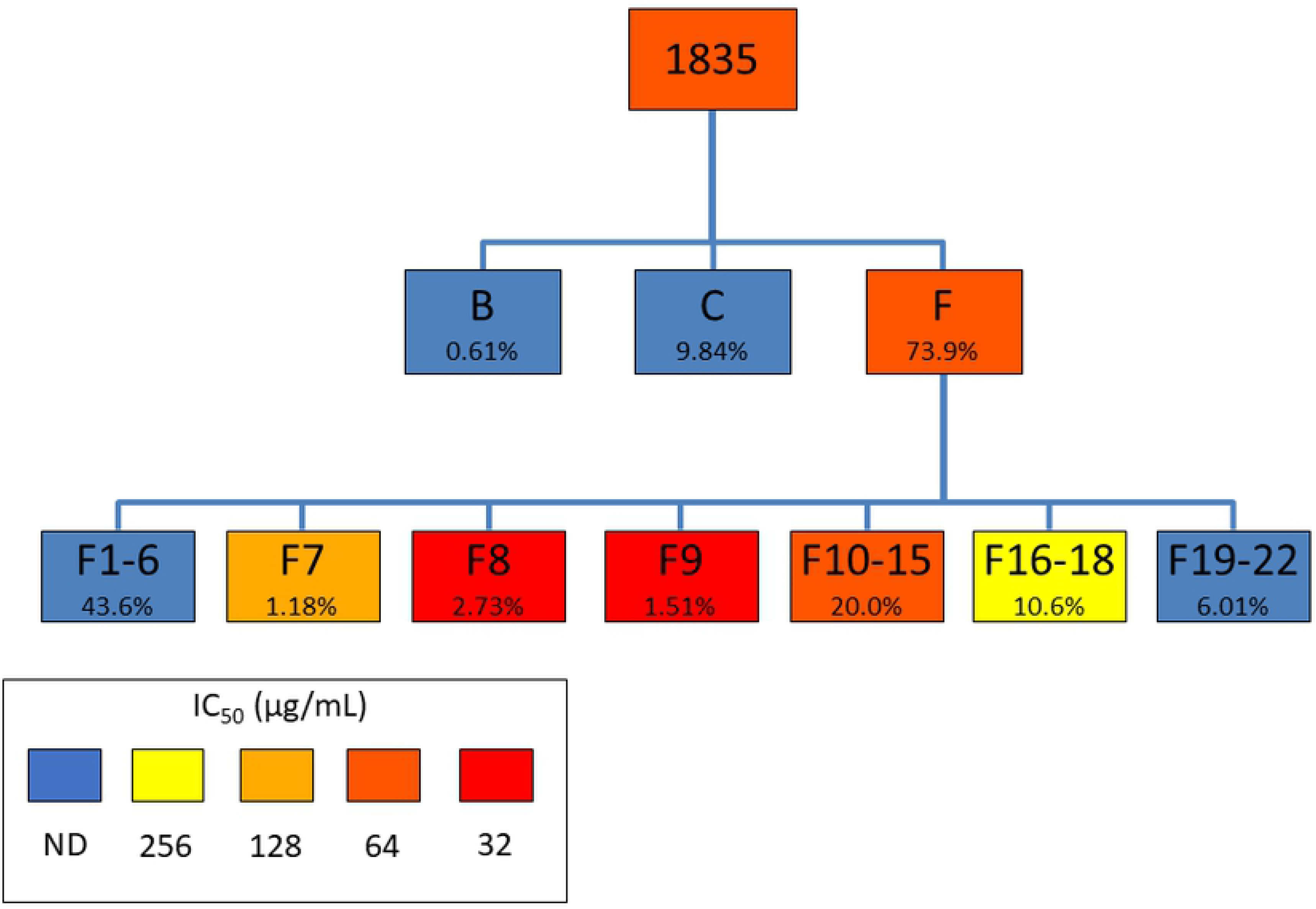
Fractionation scheme of extract 1835 (*Lechea mucronata*) showing percent yield and growth inhibition IC_50_ against *Acinetobacter baumannii*. Percent yields are relative to immediate parent.

Extract 429 (*S. terebinthifolia*) was partitioned using a modified Kupchan liquid-liquid partitioning scheme. Extraction solvents used were: hexanes, ethyl acetate, n-butanol, and water and were labeled B, C, D, and E according to solvent, respectively. The ethyl acetate partition 429C was fractionated (Fig 2) using a 330 g silica column (RediSep, Teledyne ISCO) via normal phase flash chromatography (Combi Flash Rf+ Lumen, Teledyne ISCO) utilizing a hexane:ethyl acetate gradient. A total of 9 fractions were collected with the most active fraction being 429C-F8. All subsequent preparative high-performance liquid chromatography (Prep-HPLC) were carried out using an Agilent Technologies 1260 Infinity II LC System (CA, USA) equipped with an Agilent Technologies 1200 Infinity Series Diode Array Detector detecting at 214 nm and 254nm. The column used for all subsequent Prep-HPLC purifications was an Eclipse XDB-C18 5 μm pore, 30 x 250 mm reversed phase column (Agilent). Fraction 429C-F8 was fractionated further via Prep-HPLC starting from a 98:2 mixture of a mobile phase of 0.1% (vol/vol) formic acid in water (A) and 0.1% (vol/vol) formic acid in acetonitrile (B) at a flow rate of 42.5 mL/minute and monitored for 75.5 mins. A total of 15 fractions were collected with the most bioactive fraction being 429C-F8-PF11. Fraction 429C-F8-PF11 was fractionated further via Prep-HPLC starting from a 98:2 mixture of 85:15 using a mobile phase of 0.1% (vol/vol) formic acid in water (A) and 0.1% (vol/vol) formic acid in methanol (C) at a flow rate of 42.5 mL/minute and monitored for 47 mins. A total of 14 fractions were collected with the most bioactive fraction being fraction 429C-F8-PF11-SF4.

**Figure 2.**
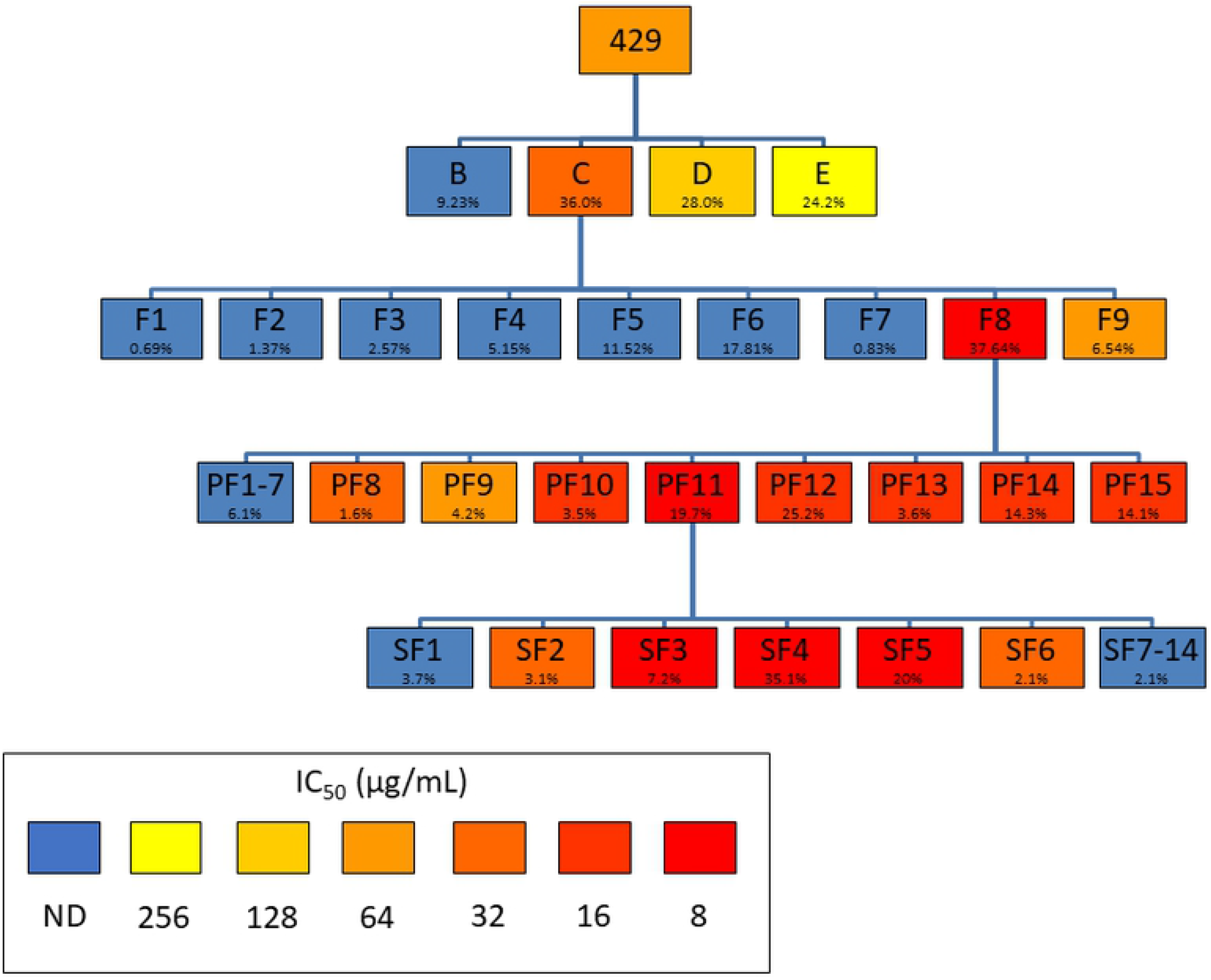
Fractionation scheme of extract 429 (*Schinus terebinthifolia*) showing percent yield and growth inhibition IC_50_ against *Acinetobacter baumannii*. Percent yields are relative to immediate parent.

*Acinetobacter baumannii* AR Bank #0035 was obtained from the CDC & FDA Antibiotic Resistance Isolate Bank (20), maintained on tryptic soy agar (TSA), and grown in cation-adjusted Mueller-Hinton broth (CAMHB) for experiments. Extract 1835, extract 429, and each partition and fraction were tested in a two-fold serial dilution gradient from 8 to 256 μg/mL (or 2 to 256 μg/mL if needed) for growth inhibition of *A. baumannii* according to CLSI methods for broth microdilution (21). Gentamicin was used as a positive control. The minimum inhibitory concentration (MIC) and the IC_50_ were defined as the lowest concentrations inhibiting > 90% and > 50% growth, respectively, relative to the vehicle control (DMSO, ≤ 2.56% of well volume).

### Calculation and interpretation of FICI

Using Loewe additivity as the basis of non-interaction, interactions can be calculated using the equation 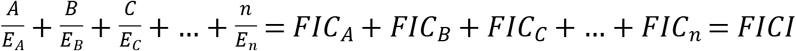, where E_A_ is the concentration of drug A alone that produces an effect (for example, 50% inhibition), A is the concentration of drug A in a mixture with the same effect, FIC_A_ is the fractional inhibitory concentration of drug A 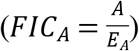, and FICI is the fractional inhibitory concentration index, the sum of the FICs of each drug present in the mixture (16). Notably, these calculations can be done with any measurable biological effect and any number of drugs in a mixture (22); for example, an investigation of the minimum inhibitory concentration (MIC) of a pairwise combination would use the form 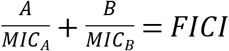.

The numerical value of the FICI is used to judge the interactions present in the mixture. An FICI value of 1 indicates no deviation from the baseline of additivity. Beyond this, interpretation of FICI has historically been debated (23, 24). Theoretically, an FICI < 1 indicates synergy and FICI > 1 indicates antagonism, but due to the variation inherent in actual experiments, FICIs of non-interacting mixtures (such as sham combinations) inhabit a range of values around 1. One widely accepted guideline for interpretation of FICI states that FICI ≤ 0.5 indicates synergy, 0.5 < FICI < 4 indicates non-interaction, and FICI ≥ 4 indicates antagonism (23). This guideline will be used for the rest of this paper.

### Calculation of EFICI

We refer to the previously undescribed application of FICI analysis to fractionated extracts as the extract FICI, or EFICI. The information needed to calculate an EFICI in any given stage of bioassay-guided fractionation is the activity of the parent extract (*P*), the yield of each fraction (*Y_n_*), and the activity of each fraction (*E_n_*). With the equation 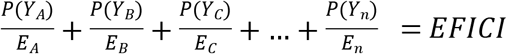, each term is one fraction. The numerator of each term – the concentration of drug/fraction in mixture that produces an effect – is calculated by multiplying the yield of the fraction (*Y_n_*) with the concentration of the parent extract that achieves the desired effect (*P*). The denominator of each term is the concentration of the fraction alone that achieves the desired effect (*E_n_*). For example, if fraction A has an MIC of 64 μg/mL and a yield of 0.2 (20%) from extract Z, which has an MIC of 256 μg/mL, 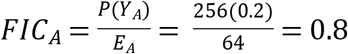. Once the FICs are calculated for each fraction that makes up a parent, they can be summed to produce the EFICI.

Using this technique, interactions can be calculated for any stage of bioassay-guided fractionation, using data from fractions and their direct parent, or interactions can be calculated for the extract as a whole, using any set of fractions that makes up the entire original extract. To assess fractions of any level that make up a parent, yield values used in calculation of FIC values for each fraction are relative to that parent. ‘Parent’ here refers to any mixture that is separated; the parent can be a crude extract or a more refined extract. For example, if extract Z is fractionated to produce fractions A, B, and C, and fraction A is itself split into subfractions *p*, *q*, *r*, and *s*, interactions can be assessed for *p*, *q*, *r*, and *s* in terms of A as a parent, or for *p*, *q*, *r*, *s*, B, and C in terms of Z as a parent. If an EFICI ≤ 0.5 is found, the indication is that the fractions interact synergistically when combined in the parent mixture.

If a fraction alone does not achieve the selected effect at any concentration tested, it can be treated as an inert substance since the term 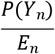 becomes vanishingly small as E*_n_*, the effective concentration, becomes arbitrarily large. Practically speaking, to calculate an FIC value, E*_n_* may be given the next highest value than the highest concentration tested (25), e.g. 512 μg/mL if the test range is a two-fold serial dilution from 2 to 256 μg/mL. Alternatively, an appropriate curve may be fitted to the data and E*_n_* may be determined by extrapolation, and FIC_*n*_ may be assigned a value of 0 if the model does not converge.

### Validation and exploration of fraction interactions

When synergy or antagonism between fractions is found, steps may be taken to validate and further investigate the interaction. First, to rule out the possibility that the apparent interaction is due to compounds lost in the process of fractionation, the fractions can be combined in a ratio based on their yields to recreate the parent extract (26). If this recreated parent has the same bioactivity as the original parent, the interaction between fractions is supported.

Determining which fractions are responsible for an interaction is more laborious, since any interaction may be due to only two fractions, more, or even all fractions that make up the parent (27). A full understanding of every interaction among the fractions of a parent, then, requires testing every possible combination of two or more fractions. For *n* fractions, this is 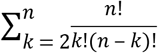, or 2^*n*^ – *n* – 1 combinations, which is 11 combinations for 4 fractions, but 1,013 combinations for 10 fractions; this approach quickly becomes impractical. However, disregarding higher-order interactions and testing only pairwise interactions requires only 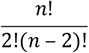, combinations, i.e. 6 combinations for 4 fractions and 45 combinations for 10 fractions. It is up to the researcher to judge on a case-by-case basis how much effort is justified in the exploration of fraction combinations.

### Accounting for experimental error

Characterization of drug interactions is an analysis one step removed from experimental data, so care should be taken to avoid the propagation of errors. In general, erring on the side of caution (i.e. assuming non-interaction when ambiguity is present) will allow for more reliable conclusions. This principle is seen in Odds’ interpretation of synergy as FICI ≤ 0.5 due to the potential 3-dilution range of any given MIC result (23).

A major example of erring on the side of caution appears in the analysis of yields of fractions. In theory, the mass of fractions collected from a chemical separation would add up exactly to the mass of the parent that was separated. The fact that this does not happen in reality (8) presents an obstacle for any analysis involving yield. When yield is used to describe how much of a fraction may be separated from a given amount of parent, it is most reasonable to calculate it using the equation 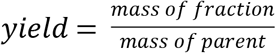. However, if yield is used to describe the relative amount of a fraction present in a mixture of fractions, it may be better to say 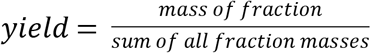. This is erring on the side of caution; for the purposes of EFICI calculation, the first equation assumes that any substance not present in the isolated fractions is completely inert, while the second equation assumes that any substance not present in the isolated fractions is a representative sample of the isolated fractions. In this paper, then, the first equation for yield is used in Figures 1 and 2 and the second equation is used in Tables 1 and 2. The closer the sum of all fraction masses is to the mass of the parent, the more precise the EFICI analysis is; this is a limitation of the method described in this paper. The recombination of fractions described in the previous section serves the purpose of testing assumptions about compounds lost in fractionation.

**Table 1.** Yields and *Acinetobacter baumannii* growth inhibition by extract 1835 and fractions.

| Fraction | Yield | IC<sub>50</sub> | FIC |
| --- | --- | --- | --- |
| 1835 | 1 | 64 | 1 |
| 1835B | 0.0072 | - | 0.0009 |
| 1835C | 0.1166 | 256 | 0.0292 |
| 1835F | 0.8762 | 64 | 0.8762 |
| 1835F-F1 | 0.4020 | - | 0.0502 |
| 1835F-F2 | 0.0133 | - | 0.0017 |
| 1835F-F3 | 0.0082 | - | 0.0010 |
| 1835F-F4 | 0.0069 | - | 0.0009 |
| 1835F-F5 | 0.0073 | - | 0.0009 |
| 1835F-F6 | 0.0086 | - | 0.0011 |
| 1835F-F7 | 0.0120 | 128 | 0.0060 |
| 1835F-F8 | 0.0279 | 32 | 0.0559 |
| 1835F-F9 | 0.0155 | 32 | 0.0310 |
| 1835F-F10 | 0.0129 | 64 | 0.0129 |
| 1835F-F11 | 0.0155 | 64 | 0.0155 |
| 1835F-F12 | 0.0525 | 64 | 0.0525 |
| 1835F-F13 | 0.0228 | 64 | 0.0228 |
| 1835F-F14 | 0.0262 | 64 | 0.0262 |
| 1835F-F15 | 0.0748 | 64 | 0.0748 |
| 1835F-F16 | 0.0563 | 256 | 0.0141 |
| 1835F-F17 | 0.0314 | 256 | 0.0078 |
| 1835F-F18 | 0.0206 | 256 | 0.0052 |
| 1835F-F19 | 0.0133 | - | 0.0017 |
| 1835F-F20 | 0.0279 | - | 0.0035 |
| 1835F-F21 | 0.0125 | - | 0.0016 |
| 1835F-F22 | 0.0077 | - | 0.0009 |

Yield calculated relative to crude extract; IC<sub>50</sub> values are µg/mL and ‘-’ indicates ‘not detected’,
and is treated as 512 for purposes of FIC calculation.

**Table 2.** Yields and *Acinetobacter baumannii* growth inhibition by extract 429 and fractions.

| Fraction | Yield | IC<sub>50</sub> | FIC |
| --- | --- | --- | --- |
| 429 | 1 | 64 | 1 |
| 429B | 0.0947 | - | 0.0118 |
| 429C | 0.3694 | 32 | 0.7387 |
| 429D | 0.2877 | 128 | 0.1438 |
| 429E | 0.2482 | 256 | 0.0621 |
| 429C-F1 | 0.0030 | - | 0.0004 |
| 429C-F2 | 0.0060 | - | 0.0008 |
| 429C-F3 | 0.0113 | - | 0.0014 |
| 429C-F4 | 0.0226 | - | 0.0028 |
| 429C-F5 | 0.0506 | - | 0.0063 |
| 429C-F6 | 0.0782 | - | 0.0098 |
| 429C-F7 | 0.0037 | 256 | 0.0009 |
| 429C-F8 | 0.1652 | 8 | 1.3731 |
| 429C-F9 | 0.0287 | 64 | 0.0287 |
| 429C-F8-PF1-3 | 0.0017 | - | 0.0002 |
| 429C-F8-PF4 | 0.0008 | - | 0.0001 |
| 429C-F8-PF5 | 0.0015 | - | 0.0002 |
| 429C-F8-PF6 | 0.0061 | - | 0.0008 |
| 429C-F8-PF7 | 0.0011 | - | 0.0001 |
| 429C-F8-PF8 | 0.0029 | 32 | 0.0059 |
| 429C-F8-PF9 | 0.0076 | 64 | 0.0076 |
| 429C-F8-PF10 | 0.0064 | 16 | 0.0255 |
| 429C-F8-PF11 | 0.0362 | 8 | 0.2896 |
| 429C-F8-PF12a | 0.0271 | 16 | 0.1082 |
| 429C-F8-PF12b | 0.0193 | 16 | 0.0770 |
| 429C-F8-PF13 | 0.0066 | 16 | 0.0263 |
| 429C-F8-PF14 | 0.0262 | 16 | 0.1050 |
| 429C-F8-PF15 | 0.0217 | 16 | 0.0869 |
| 429C-F8-PF11-SF1 | 0.0015 | - | 0.0002 |
| 429C-F8-PF11-SF2 | 0.0012 | 32 | 0.0024 |
| 429C-F8-PF11-SF3 | 0.0028 | 8 | 0.0224 |
| 429C-F8-PF11-SF4 | 0.0138 | 8 | 0.1100 |
| 429C-F8-PF11-SF5 | 0.0078 | 8 | 0.0625 |
| 429C-F8-PF11-SF6 | 0.0008 | 32 | 0.0017 |
| 429C-F8-PF11-SF7 | 0.0015 | - | 0.0002 |
| 429C-F8-PF11-SF8 | 0.0011 | - | 0.0001 |
| 429C-F8-PF11-SF9 | 0.0014 | - | 0.0002 |
| 429C-F8-PF11-SF10 | 0.0012 | - | 0.0002 |
| 429C-F8-PF11-SF11 | 0.0008 | - | 0.0001 |
| 429C-F8-PF11-SF12 | 0.0006 | - | 0.0001 |
| 429C-F8-PF11-SF13 | 0.0008 | - | 0.0001 |
| 429C-F8-PF11-SF14 | 0.0008 | - | 0.0001 |

Yield calculated relative to crude extract; IC<sub>50</sub> values are µg/mL and ‘-’ indicates ‘not detected’, and is treated as 512 for purposes of FIC calculation.

Analysis of bioactivity, the other side of EFICI calculation, raises other questions of experimental error. Is it better to describe activity using only the drug concentrations tested in experiments, or should curves be fitted to the data to allow for both extrapolation and a more continuous scale for effective concentration values? More complete discussions of this topic are found in other papers (28, 29), but for the purposes of EFICI calculation, both approaches have benefits and drawbacks. Using discrete, experimentally-verified effective concentrations is simpler and is broadly accepted for reporting bioactivity, but the lack of a continuous scale introduces error when the effective concentration values themselves are used in further analysis. Fitting curves to data may minimize the error produced by analyzing effective concentration values, but adds another layer of complexity, and improperly modeled curves (e.g. produced by irregularly distributed data or non-optimal equations or constraints) may introduce even more error than using the original discrete data. One further benefit of fitting curves to data is that it allows for more complex analysis (29) and visualization of drug interactions, such as response-surface modeling; these approaches may be advantageous in cases—such as differing relative potencies and maximal effects—in which basic Loewe additivity is an insufficient null model (10).

## Results

### Synergy in *Acinetobacter baumannii* growth inhibition by *Lechea mucronata*

Yields and individual activities of *L. mucronata* extract 1835 and its fractions are shown in Table 1. The IC_50_ values not detected in the tested concentration range (8 to 256 μg/mL) were treated as 512 μg/mL for the purposes of FIC calculation. Note that when the IC_50_ of a fraction is equal to the IC_50_ of the parent, the FIC is equal to the yield.

The EFICI of the first round of separation (1835B + 1835C + 1835F = 1835) is 0.9063. The EFICI of the second round of separation, using the 1835F-Fs in terms of 1835F, is 0.4429; for this calculation, it should be noted that since Table 1 lists FIC values in terms of 1835, the Table 1 FIC values of fractions that make up 1835F should be corrected by dividing by the FIC value of 1835F, the parent in this case. The overall EFICI of all the fractions produced (1835B + 1835C + 1835F-Fs = 1835) is 0.4181.

### Non-interaction in *Acinetobacter baumannii* growth inhibition by *Schinus terebinthifolia*

Yields and individual activities of *S. terebinthifolia* extract 429 and its fractions are shown in Table 2. IC_50_ values not detected in the tested concentration range (2 to 256 μg/mL) were treated as 512 μg/mL for the purposes of FIC calculation.

The overall EFICI of all the fractions produced (429B + 429D + 429E + 429C-F1-7,9 + 429C-F8-PF1-10,12-15 + 429C-F8-PF11-SFs = 429) is 0.9129.

## Discussion

Analysis of the example data sets showed one case of synergy in a plant extract and one case of non-interaction, both for growth inhibition of *A. baumannii*. Using the guidelines of FICI ≤ 0.5 for synergy, 0.5 < FICI < 4 for non-interaction, and FICI ≥ 4, the *Lechea mucronata* 1835 partitions are non-interactive (EFICI 0.9063), but the fractions of 1835F are synergistic (EFICI 0.4429) in terms of 1835F. The practical implication is that 1835F does not depend on other partitions for activity, but further separation of 1835F results in a loss of activity; a quantitative decision can be made to restrict further synergy studies to the fractions of 1835F. Moreover, development of 1835F as a botanical drug is supported (but not necessarily prescribed) by this analysis.

The bioassay-guided fractionation of *S. terebinthifolia* extract 429 does not exhibit a synergistic EFICI at any step of separation; partitioning and fractionation of 429C, 429C-F8, and 429C-F8-PF11 produce EFICI values of 0.9565, 1.8587, 0.5548, and 0.6915, respectively, for an overall non-interactive EFICI of 0.9129. While the most active fraction, 429C-F8-PF11-SF4, has an FIC value of 0.1100 and is therefore not responsible for all or even the majority of activity in the crude extract, it can be postulated that 429C-F8-PF11-SF4 is as active alone as it is in combination.

Since the two data types needed for this analysis are yields and effective concentrations, there are a variety of ways to prepare the relevant data for calculations, as described in the methods section and demonstrated in the results section. The comparative benefits of relative or absolute yield and discrete or modeled data are subjects for further discussion, but all these approaches have theoretical validity in the method described here. Additionally, the EFICI values produced may be subjected to any valid FICI interpretation, potentially depending on the field of study in question (23). Admittedly, reporting EFICI values alone is reductionist, but most exhaustive representations of data become impractical beyond pairwise combinations and especially difficult with the complexity of bioassay-guided fractionation. For example, fully displaying the interactions present in extract 429 would require a 38-dimensional visualization.

## Conclusions

The usefulness of interaction analysis in bioassay-guided fractionation is threefold. First, it can alert researchers to interactions present in natural substances without requiring additional experiments. Second, it can be used as a quantitative guide to synergy studies in complex extracts, yielding definitive cut-off values and supplementing the qualitative eyeballing of data when making go/no-go decisions. Third, synergistic EFICI values can give support to development of extracts or refined fractions as botanical drugs (e.g. according the FDA Botanical Drug Guidance).

Care must be taken not to over-interpret the EFICI values produced by this analysis, but with proper data management, much useful information can be gleaned about the interactions present in complex mixtures. As more combination therapies are identified and developed, this method and others like it will help give rigor to the study of drug mixtures that naturally exist in plants and other organisms.

## Declarations

### Availability of data and materials

The datasets analyzed in this study are available from the corresponding author upon reasonable request.

### Author contributions

Conceptualization: MD and CQ; Investigation: MD, LM, and MB; Formal Analysis: MD; Supervision: CQ; Writing – Original Draft Preparation: MD; Writing – Review & Editing: all authors

## Acknowledgements

Thanks to Dr. François Chassagne for commenting on the manuscript. We thank Dr. Kier Klepzig and the Jones Center at Ichauway for field research support in the collection of *L. mucronata*.

